# Riparian forest restoration as sources of biodiversity and ecosystem functions in anthropogenic landscapes

**DOI:** 10.1101/2021.09.08.459375

**Authors:** Y Antonini, MV Beirão, FV Costa, CSv Azevedo, MM Wojakowski, AR Kozovits, MRS Pires, HC de Sousa, MCTB Messias, MA Fujaco, MGP Leite, JP Vidigal, GF Monteiro, R Dirzo

## Abstract

1. Restoration of tropical riparian forests is challenging, since these ecosystems are the most diverse, dynamic, and complex physical and biological terrestrial habitats. This study tested whether biodiversity can predict ecosystem functions in a human-impacted tropical riparian forest.
2. We explored the effects of several biodiversity components (taxonomic or functional groups) on different ecosystem functions associated with restored riparian forests
3. Overall, 49% of the biodiversity components showed positive effects on ecosystem functions, each component to a different degree. In general, our results showed that both taxonomic and functional biodiversity had strong effects on ecosystem functions indicating that floral and faunal biodiversity enhanced the multifunctionality of these restored riparian tropical forests.
4. These findings indicate that in restored riparian forests, recovery of biodiversity is followed by improvement in important ecosystem functions that are the basis for successful restoration. Future research and policy for restoration programs must focus on restoring elementary faunal and floral components of biodiversity in order to promote ecosystem multifunctionality.

## Introduction

The process of habitat loss and fragmentation is the main driver of the current worldwide decline in biodiversity (Fahrig, 2003). Alterations in biodiversity and ecosystem services, largely driven by global environmental change, are also contributing to this decline (Foley et al., 2005). As a result, the number and persistence of many species will depend not only on habitat protection but also on habitat restoration, defined as the process of facilitating recovery of ecosystems following disturbance (Pedrini et al., 2020).

Tropical forests have many unique properties related to their high rates of primary productivity and biodiversity, which distinguish them ecologically from other ecosystems worldwide (Brockerhoff et al., 2017). These properties include the development of biological structures in vertical and horizontal layers of living and dead plants, a complex process at multiple vertical levels, the ability for self-renewal in the face of constant land-use changes and anthropogenic disturbances, restoring ecological functions (Martins et al 2017). These forests are comprised of multiple ecological functions that are driven by variable environmental conditions and operate at multiple spatial scales (Gardner et al., 2009). For instance, patches of forest, especially riparian forests, have a strong influence on micro- and regional climates (Allen, 2016; Burdon et al., 2020).

Tropical riparian forests are among the most diverse, dynamic, and complex biophysical habitats in terrestrial environments (Burdon et al., 2020). As interfaces between terrestrial and aquatic systems, they encompass sharp environmental gradients, complex ecological processes, and unique communities (Little et al., 2015; Pollock & Beechie, 2014). Riparian forests are recognized as important sources of “ecosystem services”, as they support watershed protection, wildlife enhancement, and ecosystem maintenance (Surasinghe & Baldwin, 2015). These forests usually support higher biodiversity and structural complexity than their surroundings (Bunnell & Houde, 2010). Consequently, deforestation of riparian areas may cause a significant decay in habitat quality in adjacent ecosystems (Surasinghe & Baldwin, 2015). Additionally, re-establishment of disturbed riparian forests is currently considered the “best management practice” for restoring aquatic ecosystems to their natural or semi-natural states (Sweeney et al., 2002).

Natural terrestrial ecosystems are valued for their ability to simultaneously maintain multiple functions and services, i.e., ecosystem multifunctionality (Allan et al., 2013). Biodiversity is by no means the only, or even the primary driver of ecosystem functioning, which is also influenced by many biotic and abiotic environmental factors that operate at different scales (Cardinale et al., 2011), but maintenance of biodiversity is a fundamental strategy for enhancing ecosystem services (Cardinale et al., 2011). For this reason, it is essential to understand how biodiversity affects different ecological processes and ecosystem functions in order to successfully restore patches of disturbed habitats (Allan et al., 2013). The relationship between biodiversity and ecosystem functioning (hereafter BEF) has emerged as one of the most exciting and controversial research areas in ecology over the last two decades (see Manning et al., 2018 for a review). Faced with the prospect of a massive and irreversible loss of biodiversity, ecologists have begun to investigate the potential consequences of current land-use changes on biodiversity and the functioning of natural and managed-novel ecosystems (Loreau et al., 2002). Biodiversity can substantially alter the structure and functioning of ecosystems and BEF studies has suggested that biodiversity loss may impair the functioning of natural ecosystems, diminishing the number and quality of services they provide (Balvanera et al. 2013; Cardinale et al. 2006, Cardinale et al. 2011, Hooper et al. 2012).

While research in the last few decades has provided many insights into BEF relationships, our current understanding of how biodiversity loss influences ecosystem functions and services amid myriad anthropogenic disturbances is neither precise nor complete (Cardinale et al., 2012; Hooper et al., 2012; Naeem et al 2012). To extend the BEF theory to restoration, researchers must gather data on ecological attributes that are easy to obtain, cost effective, and easily applicable, such as land use and canopy height, usually used to evaluate wildlife support (Palmer & Filoso, 2009). Still, no study has shown that species richness of planted trees directly increases long-term functional benefits in ecologically restored riparian forest sites (i.e., without weeding and replanting). As restored plant communities mature, their BEF relationships could be affected by trait-based changes in composition and abundance that cannot be evaluated in short-term experiments. Thus, to evaluate the success of forest restoration projects, understanding the long-term relationships between BEF is essential, insofar as it affects the ability of ecosystems to simultaneously provide multiple functions and services, in other words, the ecosystem multifunctionality (Hector & Bagchi, 2007).

To assess whether BEF analyzed at different scales (taxonomic biodiversity, functional biodiversity) might predict ecosystem multifunctionality (decomposition; leaf and miscellaneous litter production; nitrogen and phosphorus content in the litter; pH and phosphorus content of the soil; and litter and soil fertility), we studied restored fragments of tropical riparian forest, within a highly heterogeneous landscape. We tested the effect of (1) animal and plant species richness, abundance, and diversity (taxonomic biodiversity level); (2) richness and abundance of functional groups (functional biodiversity level).

## Methods

### 2.1 Study sites and restoration overview

The study was conducted in five patches of riparian forest that represent a chronosequence of restoration. The patches are in different areas (hereafter referred as sites) surrounding the reservoir of the Volta Grande hydroelectric power plant on the Rio Grande River in southeastern Brazil (20°01′54″ S, 48°13′17″ W) (Figure S1, Table S1). The region has a tropical climate with dry winters and rainy summers – classified as AW, following Köppen (Alvares et al., 2013), with a well-defined dry season between May and October and a rainy season from November to April. The mean annual temperature ranges from 22 °C to 24 °C and the mean annual precipitation reaches 1,500 mm.

The study sites are in a highly anthropogenic matrix formed mainly by grassland and sugarcane plantations. Four of the five sites have been reforested and have different ages (10 and 20 years) and widths (30 and 100 m), and the fifty site is a 30-year-old, 400-m-wide and naturally restored secondary forest, here considered as a reference site (Tables S1). Most of the original riparian vegetation in the study area was removed and flooded during the construction of the reservoir in 1974. Between 1994 and 2004, 10-month-old nursery-grown seedlings of 35 tree species, raised from seeds obtained in nearby forest remnants, were planted in a single replanting project along the shores of the reservoir, with a spacing of 3 × 2 m.

### 2.2 Experimental design

At each of the five sites, we installed four randomly plots, each 1600 m2. Biodiversity and environmental samplings were performed monthly between March 2013 and January 2014. Details of sampling methods for tree species, vertebrates (birds, small mammals, amphibians and reptiles, invertebrates (hymenoptera and soil fauna), and ecological processes can be found in the Supplementary Material.

The selected ecosystem functions, all of which are important for ecosystem multifunctionality (Maes et al., 2012), included: litter (leaf and miscellaneous) production and decomposition; litter nitrogen and phosphorus concentrations; soil pH and available phosphorus; and indexes of litter-quality and soil-fertility. Details for sampling of ecosystem functions can be found in the Supporting Information.

To disentangle the effects of distinct predictors on ecosystem functions, we divided them into two levels: taxonomic biodiversity (animal and plant species richness, including seed rain; abundance; and Shannon diversity) and functional biodiversity (animal and plant functional groups), for a total of 67 variables (Figure 1).

**Figure 1.**
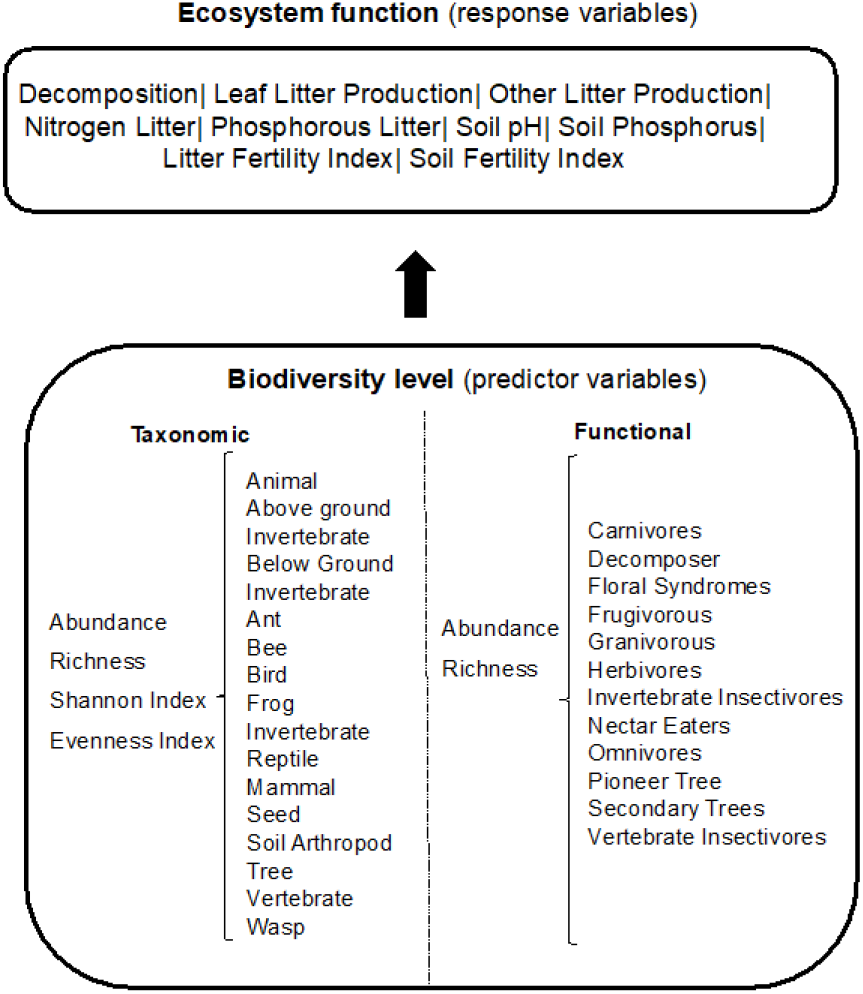
Conceptual framework indicating the level of distinct studied components: biodiversity, local, and landscape predictors from which each model was fitted to explain ecosystem functions (used as response variables).

### 2.3 Statistical Analysis

#### 2.3.1 Site dissimilarities according to land use and biodiversity

In order to understand the degree of dissimilarity of the study sites in terms of biodiversity components, we performed principal components analyses (PCA) with the package vegan for software R (R Development Team, 2016). In these analyses, sites were ordinated in relation to: (1) richness and abundance of animals and plants, (2) animal and plant diversity (Shannon and evenness indexes), and (3) richness and abundance of functional groups (for animals and plants). Prior to the PCA, we ran correlation analyses for each of the two groups (with the package pych for R) and removed the variables that were highly correlated (r > 0.8).

#### 2.3.2 Ecosystem multifunctionality analysis

To understand whether ecosystem functions can be predicted by biodiversity and environmental features, we fitted two models, structured according to different levels of sampling (Figure 1): taxonomic biodiversity model and functional components model.

Because of the large number of predictor variables, we performed a variable-selection procedure that identifies the most important variables and minimizes prediction risk, resulting in a highly interpretable model to predict forest multifunctionality under a restoration scenario. We utilized the least absolute shrinkage and selection operator analysis (lasso; Tibshirani, 1996), a shrinkage method that applies the L1 penalty to least-squares regression, thereby performing a subset selection. The amount of the penalty can be fine-tuned using a constant called lambda. First, we determined how the variables varied along the coefficients in the model; in this step, the variables that did not change were eliminated. Then, we selected the minimum lambda to obtain the mean cross-validated error and the coefficient for each variable. The lasso analysis was executed with the package glmnet for R (Friedman et al., 2010). We conducted all statistical analyses using the R programming language (R Development Team, 2016).

## Results

### 3.1 General results for biodiversity

During the sampling period, we captured 58,858 individual animals of 268 species, including 16 mammals, 122 birds, 23 amphibians and reptiles, 28 species of cavity-nesting bees and wasps, 79 species of ants, and 451 morphospecies of soil invertebrates. We sampled 127 tree species for a total of 1006 individuals. From these taxa, we classified 24 functional groups including richness and abundance of carnivores, herbivores, frugivores, granivores, invertebrate and vertebrate insectivores, decomposers, nectarivores, pioneer and secondary trees, and floral syndromes.

### 3.2 Site dissimilarities according biodiversity

Altogether, the richness and abundance of different biodiversity groups explained 69% of site dissimilarities (SFigure 1a). Biodiversity effects, on ecosystem functions, strengthened with time as a consequence of time since restoration. Overall, sites 1 (30 years old), 2 and 3 (20 and 10 yo, respectively) were related to higher richness and abundance of trees, seeds, and birds (axis 2). In contrast, sites 4 (20 yo) and 5 (10 yo) were associated with decreases in the richness and abundance of invertebrates (e.g., wasps) and vertebrates (e.g., mammals) (axis 2). Site dissimilarities according to the Shannon diversity and evenness of general groups (SFigure 1b) suggested that sites 1 and 2 are more similar to each other, while sites 3, 4, and 5 are closer to each other.

The richness and abundance of functional groups explained 69% of site dissimilarities (SFigure 1c). In general, sites 1, 2 and 3 were related to high richness and abundance of pioneer and secondary trees, frugivores, and omnivores on axis 2 (42% explanation). On the other hand, sites 4 and 5 were more similar to each other, being related to low richness and abundance of the functional groups, on both axes.

### 3.3 Ecosystem multifunctionality analysis

A total of 56 (out of 118) predictor variables influenced at least one of the nine ecosystem functions analyzed. Around 40% of were positive. This percentage of explanation varied for each ecosystem function and biodiversity levels.

The summary results for the lasso analyses are presented in Figures 2 and 3, distinguished according to the two fitted models. In these figures, the y-axis displays the lasso-selected predictor variables, and the x-axis represents the coefficient estimates for each variable. Only those coefficients with values different from 0 were displayed on the plot. A negative coefficient implies a negative effect on the response variable (i.e., the ecosystem function), and a positive coefficient, a positive effect. Below, we described each model fitted according to the biodiversity levels.

**Figure 2:**
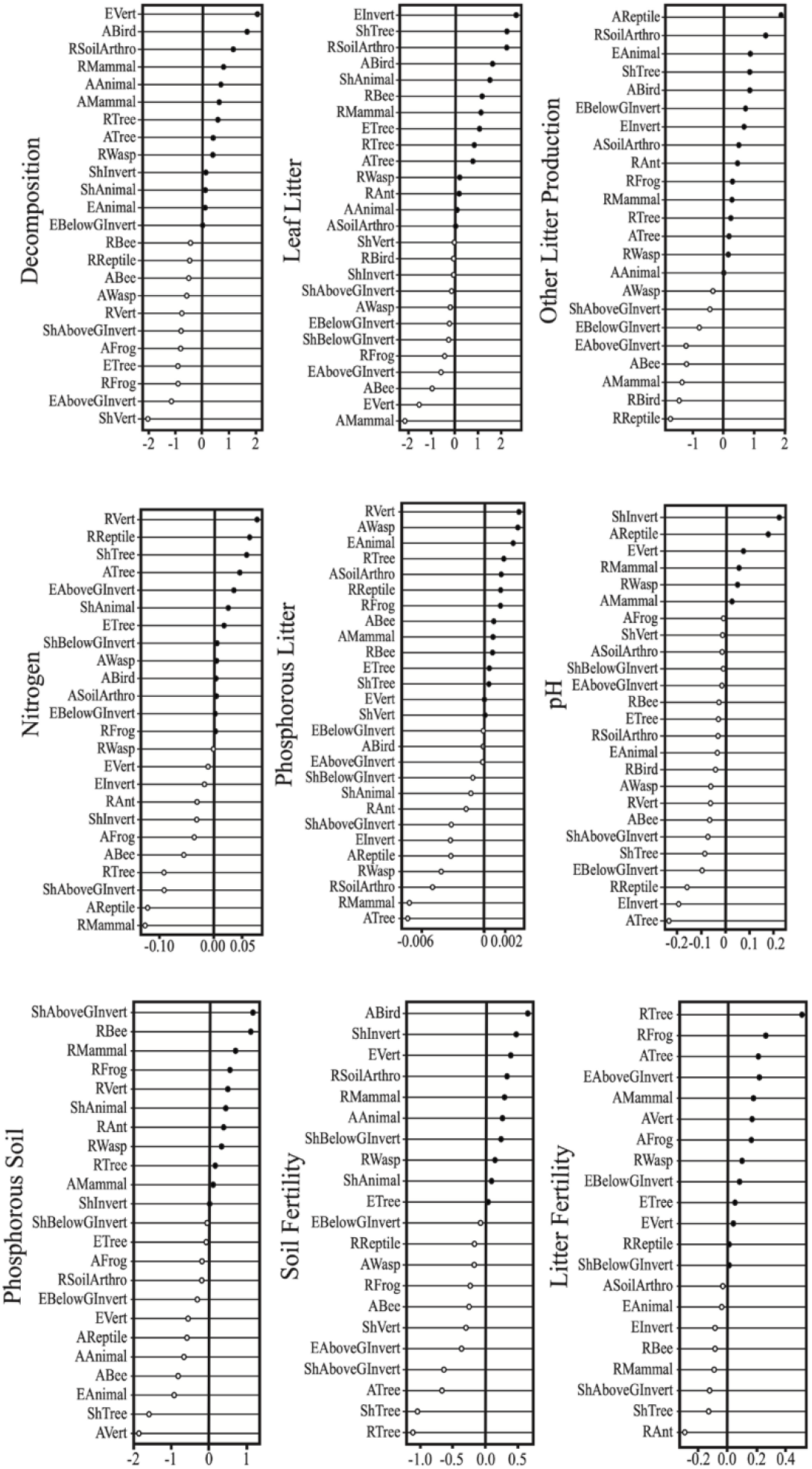
Taxonomic biodiversity components effects (taxonomic biodiversity model) on distinct ecosystem functions (decomposition, leaf in the litter, other litter production, Nitrogen, Phosphorous in the litter, pH, Phosphorous in the soil, soil fertility and litter fertility). W-axis represents β-coefficient from lasso analysis. Values greater than 0 indicate positive effect (dark circles) and lower than 0 negative effect (light circles). Central line represents 0 values.

**Figure 3:**
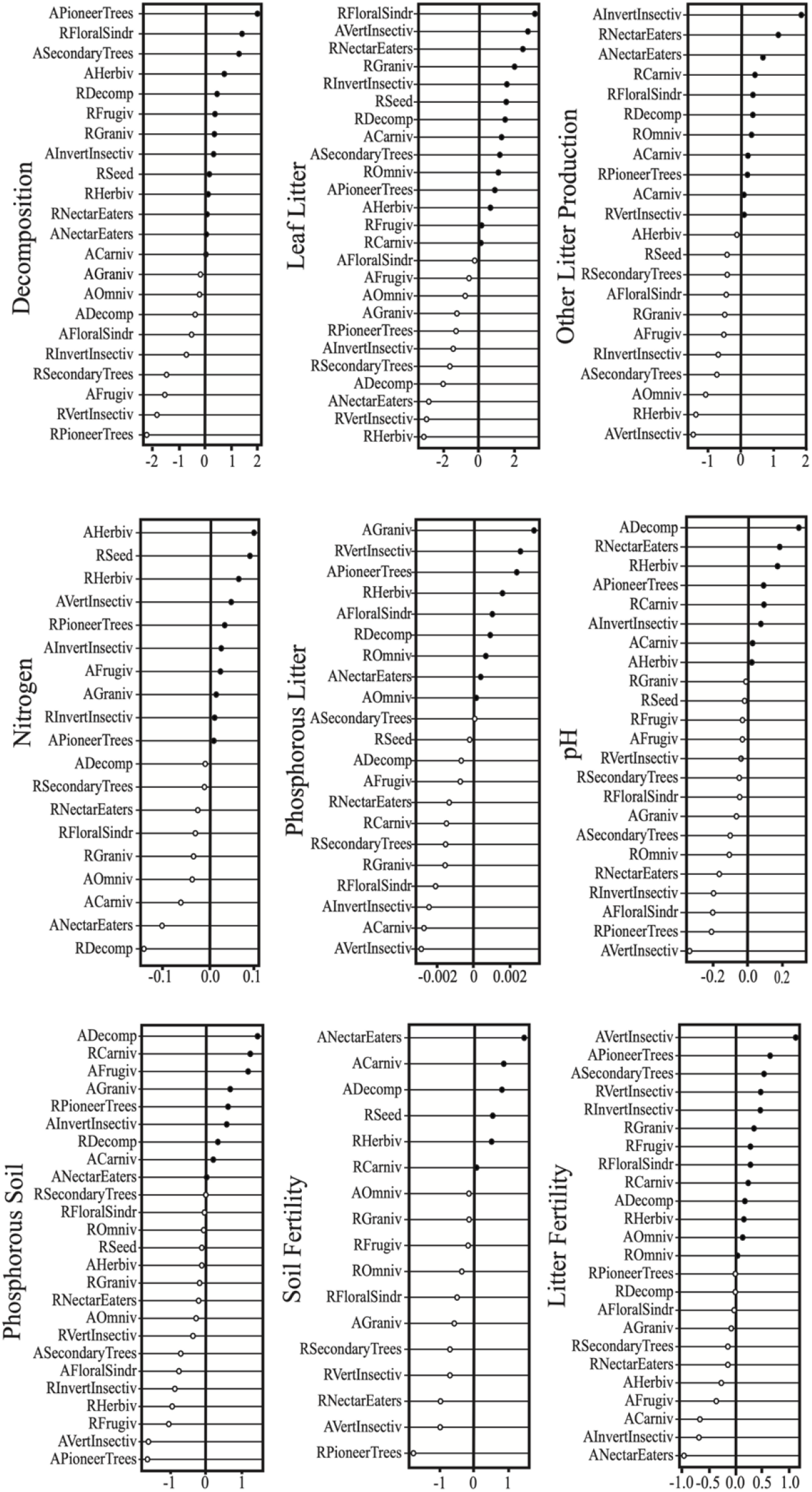
Functional Biodiversity models on the ecosystem functions (decomposition, leaf in the litter, other litter production, Nitrogen, Phosphorous in the litter, pH, Phosphorous in the soil, soil fertility and litter fertility). W-axis represents β-coefficient from lasso analysis. Values greater than 0 indicate positive effect (dark circles) and lower than 0 negative effect (light circles). Central line represents 0 values.

### 3.4 Effects of biodiversity model on ecosystem functions

In general, the species richness and abundance of distinct taxonomic groups had 50% positive effects on ecosystem functions. The taxonomic biodiversity variables with the most positive effects on the ecosystem function were richness of trees and wasps, affecting six of the nine ecosystem functions. The Shannon diversity of above-ground invertebrates negatively affected eight of the nine ecosystem-function variables.

Of the 22 predictor variables selected by lasso in the taxonomic biodiversity model, 12 had a positive effect on litter decomposition rates (Figure 2). Evenness of vertebrates, abundance of birds, and richness of soil arthropods had the three most-positive effects on decomposition. On the other hand, the abundance of bees, frogs and wasps and the richness of vertebrates (for example) had a negative effect on litter decomposition (Figure 2).

Overall, the richness and abundance of different taxa (Figure 2) showed positive effects on leaf litter and other litter production. Therefore, 55% of the predictors of taxonomic biodiversity components showed a positive effect on leaf and other litter production. On average, the Shannon diversity and evenness of trees, richness of soil invertebrates and abundance of birds had the higher positive effect on the rates of leaf and other litter production. On the other hand, some taxonomic-biodiversity predictors (e.g., richness of birds, frogs and bees, and abundance of bees) had small influences on leaf and other litter production (Figure 2).

Most of the taxonomic biodiversity predictors (13 of 24; 54%) had a positive effect on nitrogen content in leaf litter. The predictors with higher positive effects included the richness of vertebrates, reptiles and Shannon diversity of trees. In contrast, we observed very small effects of the abundance of reptiles, richness of mammals, and the Shannon diversity of below-ground invertebrates on nitrogen content in litter (Figure 2).

Additionally, we found that 14 (of 27; 52%) of the predictor variables at the taxonomic biodiversity level had positive effects on phosphorus content in litter. Among these, the richness of vertebrates, abundance of wasps and Evenness of animal showed higher positive effects. On the other hand, the abundance of trees, richness of mammals and richness of invertebrates of soil ants showed small effects on phosphorus content in litter (Figure 2).

Only six predictors of taxonomic biodiversity (of 25; 24%) had a positive effect on soil pH. The Shannon diversity of invertebrates, the abundance of reptiles, and the evenness of vertebrates had the highest positive effects on the pH. In contrast, the abundance of trees, Evenness of invertebrates and richness of reptiles, had the most negative impact (Figure 2).

For the phosphorus content in soil, 11 of 25 (44%) of the taxonomic predictors had positive effects with Shannon diversity of above ground invertebrates, richness of bees and mammals showing the highest positive effect. Otherwise, the Shannon diversity of trees, abundance of vertebrates and Evenness of animal had negative effects on the phosphorus content in the soil (Figure 2).

In general, soil fertility were affected by 10 out of 21 (48%) taxonomic-biodiversity predictor variables (Figure 2). Thus, the abundance of birds, Shannon diversity of invertebrates and Evenness of vertebrates had a higher effect soil fertility. On the other hand, richness, abundance and Shannon diversity of trees had very small effect on soil fertility (Figure2).

In general, litter fertility was affected by 13 out of 21 (62%) taxonomic-biodiversity predictor variables (Figure 2). Thus, the richness and abundance of trees and had a higher effect on litter fertility. On the other hand, Shannon diversity of above ground invertebrates and trees and richness of ants had small effects on litter fertility.

### 3.5 Functional-biodiversity effects on ecosystem functions

Of the 22 predictors of functional biodiversity selected to explain decomposition, we found that 95% had a positive effect on at least two functions (from 2 to 7 of the 9) (Figure 3).

The abundance of pioneer and secondary trees and richness of floral syndromes had higher positive effects on decomposition (Figure 3). On the other hand, abundance of frugivorous, richness of vertebrate insectivorous and richness of pioneer trees had small effects on decomposition (Figure 3).

The majority (56%) of the predictor variables at the functional biodiversity level (14 of 25) had positive effects on the leaf litter. Richness of floral syndrome, abundance of vertebrate insectivores and richness of nectar eaters, had the large positive effect on the leaf litter. On the other hand, abundance of nectar eaters, vertebrate insectivores and richenss of herbivors had small effect on leaf litter (Figure 3).

Of the 22 predictor variables at the functional biodiversity level, 11 (50%) had a positive effect on other content in litter production, although the sizes of their effects differed. For example, the abundance of vertebrate insectivores and richness and abundance of nectar eaters had a stronger effect on other litter production. Conversely, the abundance of omnivores, richness of herbivores and abundance of invertebrates insectivorous had negative effect on other litter production (Figure 3).

Many of the functional-group predictors (10 of 19-53%) had positive effects on nitrogen content in litter (Figure3). The abundance and richness of herbivores and richness of seed eaters had the highest positive effects on nitrogen content in litter. On the other hand, abundance of carnivores and nectar eaters and richness of decomposers had small effects on nitrogen in litter.

For phosphorus content in litter, nine of 21 (43%) predictors had a positive effect. Thus, the abundance of granivores, pioneer trees, richness of insectivorous vertebrates, had positive effects. In contrast, the abundance of invertebrate and vertebrate insectivores and abundance o, carnivores, among other predictors, had small effects on phosphorus in litter (Figure 3).

For pH of soil, only eight out of 22 (36%) predictor had positive effects. Increases in the abundance of decomposers and richness of nectar eaters and herbivores had higher positive effects on soil pH (Figure 3). In contrast, the richness of pioneer trees, abundance of floral syndrome and vertebrate insectivores had small effects on soil pH (Figure 3).

For phosphorus content in soil, nine of 25 (36%) predictors had a positive effect. Thus, the abundance of decomposer and frugivorous and richness of carnivores had the highest positive effects. In contrast, the richness of frugivorous and abundance of vertebrate insectivores and pioneer trees, among other predictors, had small effects on phosphorus content in soil (Figure 3).

For soil fertility only six out of 17 (35%) selected predictors had positive effect. The abundance of nectar eaters, carnivores and decomposers showed higher positive effects on soil fertility. On the other hand, richness of nectar eaters and pioneer trees and abundance of vertebrate insectivores had small effects (Figure 3).

Finally, for litter fertility 13 out of 24 (54%) predictors had positive effect. The vertebrate insectivores are one predictor of the first tree functional biodiversity having higher positive effect, together with richness and abundance of pioneer trees. On the other hand, abundance of carnivore invertebrate, insectivores and nectar eaters had higher negative effect on litter fertility (Figure 3).

## Discussion

Our results showed that biodiversity effects seem to be remarkably consistent across different groups of organisms and among trophic levels and functional groups. This consistency indicates the existence of general underlying principles that dictate how the organization of biological communities’ influences ecosystem functioning (Hooper et al., 2012). We found exceptions to this pattern for some BEF; however, there was substantial variability in the response of ecosystem functions under different environmental conditions. In general, we found that for each sampling scale, half of the predictors, on average, had strong positive effects on the ecological processes studied, while the other half caused small or null effects. The richness and abundance of different biodiversity groups were important predictors to define the similarities of ecosystem functions between our reforested study sites (e.g., tree density and the size and age of sites). The restored sites are located in areas with diverse land uses, including monocultures of sugarcane, soybean, and rubber trees, which in many cases are the dominant matrices. Previous studies in the same area, have suggested that landscape configuration has a strong effect on the local biodiversity and consequently on some ecosystem functions (Araújo et al. 2018; Londe et al. 2020). Studies including planting experiments have demonstrated that larger and older areas have experienced persistent positive diversity–productivity relationships (Tilman et al., 2006; Van Ruijven & Berendse, 2010), while smaller and younger reforested patches have commonly lost this relationship, or it is considerably weakened (Roscher et al., 2012).

### 4.1 Biodiversity effects on ecosystem functioning

Our first question was whether taxonomic biodiversity had any effect on ecosystem functioning in reforested riparian areas. We expected that an increase in biodiversity would have a positive effect on the ecosystem functions. We found that an increase in overall animal (of all the taxonomic groups sampled) and tree species richness and abundance, the diversity index, and evenness had positive effects on 54% of the ecosystem functions. For example, richness and abundance of mammals, abundance of birds, richness of arthropods in the soil, and richness of trees all had positive effects on a minimum of five and a maximum of six of the nine ecosystem functions studied. However, there were exceptions, where some biodiversity taxa had minimal or null effects (e.g., bird richness, overall vertebrate abundance, and bee abundance).

We found that tree species richness and abundance had positive effects on a large number of processes in the restored ecosystems, although they did not show any effect on nitrogen and phosphorus in litter or on soil fertility, as expected. This could be explained by the more dystrophic soil of the reference patch of forest, with the highest species richness and abundance, compared with the reforested patches (Szefer et al., 2017). There is growing evidence that the quality of leaf litter is related to the ecological role played by functional groups of species (Szefer et al., 2017) and to variation in soil N and P availability (Kozovits et al., 2007; Hobbie, 2015). Several soil physical parameters can affect the relationship between soil fertility and plants, such as the percentage of clay minerals, soil aggregate stability, and soil compaction (Bardgett et al., 2014). All these physical parameters influence soil hydrological regimes and consequently the exchange of chemical elements (Horn & Gra, 1998; Cheng & Heidari, 2019), especially P and N, which are directly related to vegetation parameters; and can affect the productivity of the ecosystem. On the other hand, soil fertility was positively affected by a few components of the biodiversity level, such as the richness of mammals, the abundance of decomposers, and the richness of seeds in seed rain.

When we evaluated the effects of biodiversity on the rate of organic-matter decomposition in litter, we found considerable variation in the predictive power of different taxonomic groups. However, as expected, the trend was more pronounced for certain taxa. For example, the abundance of birds, richness of soil arthropods, and richness and abundance of trees were positively related to the litter decomposition rate. According to Cardinale et al. (2011), limited evidence suggests that, on average, a decline in plant diversity may reduce decomposition rates and the efficiency by which biologically essential elements are recycled back into their inorganic forms. The lack of a direct and strong relationship between the tree diversity and soil processes such as decomposition may also be a result of oversimplifying the data analysis. Trees support other components of diversity in the system, such as understory herbaceous plants and soil microorganisms, among other actors that mediate the litter decomposition process. Explanatory models that include multitaxonomic diversity reveal a significant indirect effect of trees on decomposition (Fujii et al., 2017).

Another important ecosystem function, the amount of litterfall produced by forests, also could be increased by augmenting biodiversity, since litterfall has components of both plant and animal origin. However, in our study, the variables that explained litter production, at the biodiversity level, showed some unexpected results, with essential components explaining little or almost nothing of ecological processes (e.g., richness and abundance of invertebrates and trees). However, the above mentioned result is consistent with other studies (Fayle et al., 2015; Oliver et al., 2015; Huang et al., 2017), since the different groups that comprise biodiversity exhibit behaviors and participate in functional groups that relate differently to the resources offered by the forest. Therefore, some groups will be more related to leaves and other groups will be more related to branches, fruits, and seeds.

The richness of soil arthropods was another biodiversity component that was positively related to litter production. Many arthropods that nest in forest soil, such as ants, termites, and coleopterans, use the forest canopy as a substrate for foraging (Souza-Campana et al., 2017; Dambros et al., 2018). A large part of the soil fauna, in our study, was composed of leaf-cutting ants and termites, as is typical in neotropical forests (Fujii et al., 2017), which could be acting to increase the quantity and quality of some organic material in the litter. Other components of biodiversity, such as bird and bee richness, were also strongly associated with litter production. This could be an indirect result, since the richness of birds and bees is linked with forest structure, with more-structured forests supporting more bird species (Casas et al., 2016; Rhoades et al., 2018). Litter production in more-structured forests exceeds that in less-structured forests (Capellesso et al., 2016; Souza et al., 2019). This result may reflect the success of the restoration process at the sites evaluated here. Also, birds are predators of invertebrates that consume litter, and predation on these invertebrates can increase the amount of litter (Stratford & Şekercioğlu, 2015). At the study sites, various species of insectivorous birds (ground, understory, and canopy) were recorded (Mafia & de Azevedo, 2020). As expected, the abundance of plants was directly proportional to leaf litter production.

The general effect of species richness differed amongst the studied functions and biogeochemical cycles (such as phosphorus and nitrogen content). Certain taxa seemed to be more important in explaining BEF. For example, the positive relationship of mammal and tree species richness with nitrogen and phosphorus concentrations in soil was expected, but the null effect of decomposer richness on these two elements was not. Balvanera et al. (2006), in a meta-analysis of biodiversity effects on ecosystem function, did not find similar results for biogeochemical cycles, which may occur if complementarity, facilitation, and insurance effects increase the community-level use of limiting resources (Hooper et al., 2012). The presence of certain groups such as legume trees may be more determinant for nutrient cycling than is species richness (Vitousek & Howarth, 1991). However, changes in vegetation composition may cause a discrepancy between biogeochemical cycles (Pasut et al., 2020). Animal bodies, feces, and fruits processed by animals are available to become soil organic matter along with litter directly produced by plants. Also, large-bodied seed dispersers such as peccaries and primates ingest, digest, and defecate large amounts of fruit pulp and seeds, as well as grasses and leaves (Fragoso & Huffman, 2000; Stevenson & Guzmán-Caro, 2010), moving plant matter across the landscape and processing it in ways that make it available to a wider range of invertebrates, fungi, and microorganisms.

The number of taxonomic biodiversity variables related to pH and phosphorus content in the soil was smaller than expected. For pH, only the diversity of invertebrates is among those expected to affect pH, since many are decomposers. We expected that plant diversity would affect pH, but we failed to find such an effect, probably because the soil of the most mature and diverse forest is more dystrophic and acidic than the eutrophic soil in the restoration patches. This weak association of biodiversity with soil pH has been reported previously. Dawud et al., (2017) found a positive effect of diversity and a negative effect of species composition on topsoil pH. Indeed, some authors have suggested that functional groups of trees are more important than biodiversity per se (Dawud et al., 2017) and have emphasized the importance of additive effects of diversity on the abundance and community structure of soil microbial and macrofaunal communities (Scheibe et al., 2015; Wandeler et al., 2016).

Litter nutrients are important for maintaining ecological processes and are strongly related to biodiversity, as the primary and secondary decomposition of organic material and the primary productivity are dependent on plants and animals (Kerdraon et al., 2020). We found positive effects for most predictor variables of taxonomic biodiversity, such as richness and abundance of trees, for both nutrients (P and N) and soil arthropods, which were good predictors for nitrogen content in litterfall. Among the positive effects, we first discuss the role of the faunal diversity in decomposition and nutrient (N and P) content in both the litter and the soil. The overall richness and abundance of animals and trees had strong positive effects on phosphorus in soil. The principal forms of phosphorus in soils are associated with calcium (Ca) or magnesium (Mg) in phosphates (relatively unweathered environments), and with clays and iron (Fe) and aluminum (Al) oxides, in old, highly weathered tropical landscapes (Spain et al 2018). The low solubilities of these phosphates and oxides make P a relatively immobile element in its inorganic form. Thus, the concentration of exchangeable phosphorus in highly weathered P-depleted soils is determined mostly by biological recycling processes, especially those related to organic-matter degradation (Tiessen, 2015). Tree diversity, in part, is important for maintaining the nitrogen and phosphorus pools in restored tropical forest (Zeugin et al., 2010), although this relationship depends on the initial site conditions (Redondo-Brenes & Montagnini, 2006) which makes robust generalizations difficult.

Plant species richness can increase fine root biomass and length, facilitating P uptake from the different soil layers. Tree species richness also has a positive effect on soil organic carbon and litter decomposition, increasing the bioavailable P content (Wu et al., 2019). Furthermore, the amount and rate of nutrient cycling are partly affected by herbivores through litterfall dung (Fonte & Schowalter, 2005). Insect herbivores can increase soil N and P fluxes by as much as 30% in tropical rainforests, through their fragmentation activity (Schowalter et al. 2011). Defecation by monkeys and other vertebrate herbivores, with further processing by dung beetles, contributes to improving soils and ultimately affects nutrient storage in these forests (Neves et al., 2010). Soil fertility depends on nutrient mineralization, and soil organic matter increases with plant richness; the expected richness of tree species determined, in this study, the greater fertility of the soil and the amount of litter produced. The richness and abundance of other animal groups such as nectarivores also had positive effects on soil fertility.

Shannon diversity and evenness also positively affected the ecosystem functions. We found positive responses and some consistency for BEF. For example, the diversity index of the overall fauna was a good predictor of important ecosystem functions, such as decomposition, N in litter, and P in soil. Moreover, Shannon diversity of the overall animal group and below-ground animals improved different ecosystem functions by more than 50%. Although we had expected that the diversity of trees (Shannon index) would have a positive effect on litter decomposition, the effect was small. The lack of a direct and strong relationship between tree diversity and soil processes such as litter decomposition may also be a matter of oversimplifying the data analysis. Trees are important in supporting other components of diversity in the system, such as understory herbaceous plants and soil microorganisms, among other actors that mediate the litter decomposition process. When explanatory models include multitaxonomic diversity, a significant indirect effect of trees on decomposition is revealed (Fujii et al., 2017). A modeling study by (Loreau & Hector, 2001) demonstrated a negative effect of plant litter diversity on litter decomposition, as a larger number of litter types should increase the probability that decomposers will not consume at least part of them. The same model predicted a positive effect of decomposer diversity on decomposition rates, due to partitioning of resources between different decomposers. Nevertheless, we found only a small effect of below-ground invertebrates on litter decomposition, although the diversity of the overall invertebrates positively influenced it.

### 4.2 Functional-diversity effects on ecosystem multifunctionality

We found that specific functional groups of organisms were essential to maintain the functions in the restored sites. Carnivores, herbivores, and pioneer trees positively affected most of the ecosystem functions (six of nine). Likewise, decomposers, insectivorous vertebrates, and nectarivores showed a positive effect on five of the nine ecosystem functions. Numerous well-known studies have posited that species identity and biodiversity are surrogates of functional-trait effects on ecosystem functioning (see Hättenschwiler et al, 2018; Szefer et al., 2017). However, according to (Schoolmaster et al., 2020), these surrogates should not be assumed to be “causal” although significant biodiversity–ecosystem function correlations are spurious associations that arise from common-cause confounding in mis-specified trait-based ecosystem function models. Residual effects of species identity, while causally related (i.e., elements of species composition), also indicate incomplete trait information.

We observed that functional-group diversity had strong effects on certain ecosystem functions, in particular those associated with litter decomposition, litter quality, and N and P cycling. Our results agreed with several BEF experiments that have shown that functional-group diversity is a good predictor of ecosystem multifunctionality (Temperton et al., 2007; Fujii et al., 2017). For example, we found a positive effect of the richness of carnivorous and herbivorous animals on litter quality and litter-P. Most measures of nitrogen increased with the abundance of pioneer trees, since many of them are legumes, able to fix atmospheric nitrogen and therefore increase nitrogen stocks (Houlton et al., 2008; Oelmann et al., 2007; Temperton et al., 2007). Indeed, we found positive effects of the richness and abundance of pioneer trees on N and P contents in the litter, as well as a positive effect of the abundance of secondary trees. Likewise, important functional groups such as herbivores and decomposers had positive and strong effects on almost 70% of the functions. According to (Dawud et al., 2017), functional groups are important in ecosystem multifunctionality, indicating that supporting a large degree of heterogeneity in specific characteristics of some taxonomic groups (those that can be captured by functional-trait diversity) may enhance ecological functions.

Increasing evidence shows that the critical means by which species influence ecosystem functions is through their functional traits (e.g., phenotypic attributes that represent niche exploitation; (Díaz et al., 2007). While functional diversity may theoretically increase with species richness in some contexts (Hooper et al., 2005), measures of taxonomic biodiversity (particularly species richness) have proved to explain little of the variance in ecosystem functions compared to indices of functional traits. Indeed, we found that certain functional groups had stronger effects on certain BEF. These components included the presence or relative abundance of certain functional trophic groups, such as herbivores, carnivores, and pioneer trees (i.e., legumes), and also an element that encompasses a functional-trait value or importance to the BEF (e.g., pollination syndromes). These are hereafter jointly termed (variation in) functional composition.

We found important effects of certain functional groups that clearly affect litterfall production. The abundance of insectivorous and nectarivorous vertebrates and seed eaters that travel through the canopy and manipulate parts of the plants contribute to the fall of leaves, seeds, and branches. Also, the abundant herbivores such as ants, termites, and beetles have a similar role as the above functional groups. However, here, the functional groups that determined soil fertility were less abundant than the groups that determined litter quality.

For some functional groups, the effect on BEF proved to be a cross-effect, for example the richness of floral syndromes, which can be related to the richness of plants. Other functional groups, where a positive effect on BEF was expected, had negative effects, such as the abundance of decomposers and decomposition. The functional groups that were more positively related to pH and phosphorus content in soil were the abundance of decomposers (as a result of organic-matter degradation processes), the richness of pioneer trees, abundance of frugivores (manipulation of the fruits that fall to the ground, while at the same time these frugivores may defecate while eating the fruits). These functional groups were consistent and expected. Several investigators have reported correlations between soil properties, such as pH or phosphorous content, with forest properties, such as above-ground biomass or species distributions (Condit et al., 2013; Schaik & Mirmanto, 2013). Pioneer trees may grow several meters in a year, improving soil fertility by accelerating soil organic-matter accumulation, enhancing P concentration, and lowering pH (Diemont et al., 2006; Vleut et al., 2013). The abundance of frugivores (animal feces; seeds and fruits that may fall to the ground) may increase the supply of nutrients and organic-matter content to the soil, leading to more favorable soil physical and chemical conditions for environmental restoration.

## Conclusion

Our findings indicated that many important ecosystem functions were highly affected by the presence of different groups at the levels of taxonomic biodiversity and functional biodiversity, which may indicate that the community shows complementarity in functional redundancy. At present, we know little about the biological mechanisms that are responsible for complementarity among species, besides, some studies showed that species loss has adverse effects on a range of ecosystem functions and services (Balvanera et al., 2006; Cardinale et al., 2006), but that relatively few species are needed to sustain the overall health of the environment (Cardinale et al., 2006), suggesting a high degree of functional redundancy (Schoolmaster et al., 2020).

Restored patches must meet two broad conservation objectives: representativeness and persistence (Noss et al., 2012). The first objective attempts to represent the variety of populations, species, or ecosystem functions of each region; while the second attempts to promote the persistence of these elements over the long term (Margules & Pressey, 2000). Our findings indicate that the restored sites represent the natural variation in taxonomic biodiversity that was almost as important for ecosystem functioning as the natural variation in functional biodiversity, but each component display specific responses. Therefore, estimating if taxonomic biodiversity is a better predictor than functional biodiversity for ecosystem functions is worth evaluating. In our study, both these approaches were essential in explaining the ecosystem multifunctionality. However, the relative importance of taxonomic biodiversity versus functional composition depended strongly on the type of ecosystem function. Although these challenges will not be easily met, the field of BEF research now has all the necessary tools to take these next important steps.

## Acknowledgements

We would like to thank the former students Adriele Magalhaes, Antonio J. Cruz, Auria Tonaco, Gustavo Araujo, Luciana Oliveira, Matheus Correa, Paola Oliveira, Pedro Maffia for contributing data. The technical staff of CEMIG Rafael Fiorine and Luciana Magalhães. A special thanks to José Ricardo da Silveira – who planned and who planned and implemented the forest restoration. The staff of Grupo Colorado, Grupo Raizen and Fazenda Noboro for maintaining the sites and for logistic support during field experiments. This project was funded by the Fapemig/CEMIG (Grant 03055/11 to Y.A.). Version 3 of this preprint has been peer-reviewed and recommended by Peer Community In Ecology (https://doi.org/10.24072/pci.ecology.100096).

## Data, scripts and codes availability

Data and scripts are available online: https://knb.ecoinformatics.org/view/doi:10.5063/G73C37

## Ethics Statement

Trapping and handling were approved by IBAMA – Instituto Brasileiro do Meio Ambiente e dos Recursos Naturais Renováveis (permissions 10717, 129311-1, 36206-1, 36206-2, 367581, 36758-2, 37067-1) and Ethics Committee of the Institute of Sciences - University Federal de Ouro Preto (Comissão de Ética no Uso de Animais– CEUA - http://www.ceua.ufop.br) (055/2012 and 142/2013).

## Supplementary information

**Supplementary Figure 1:**
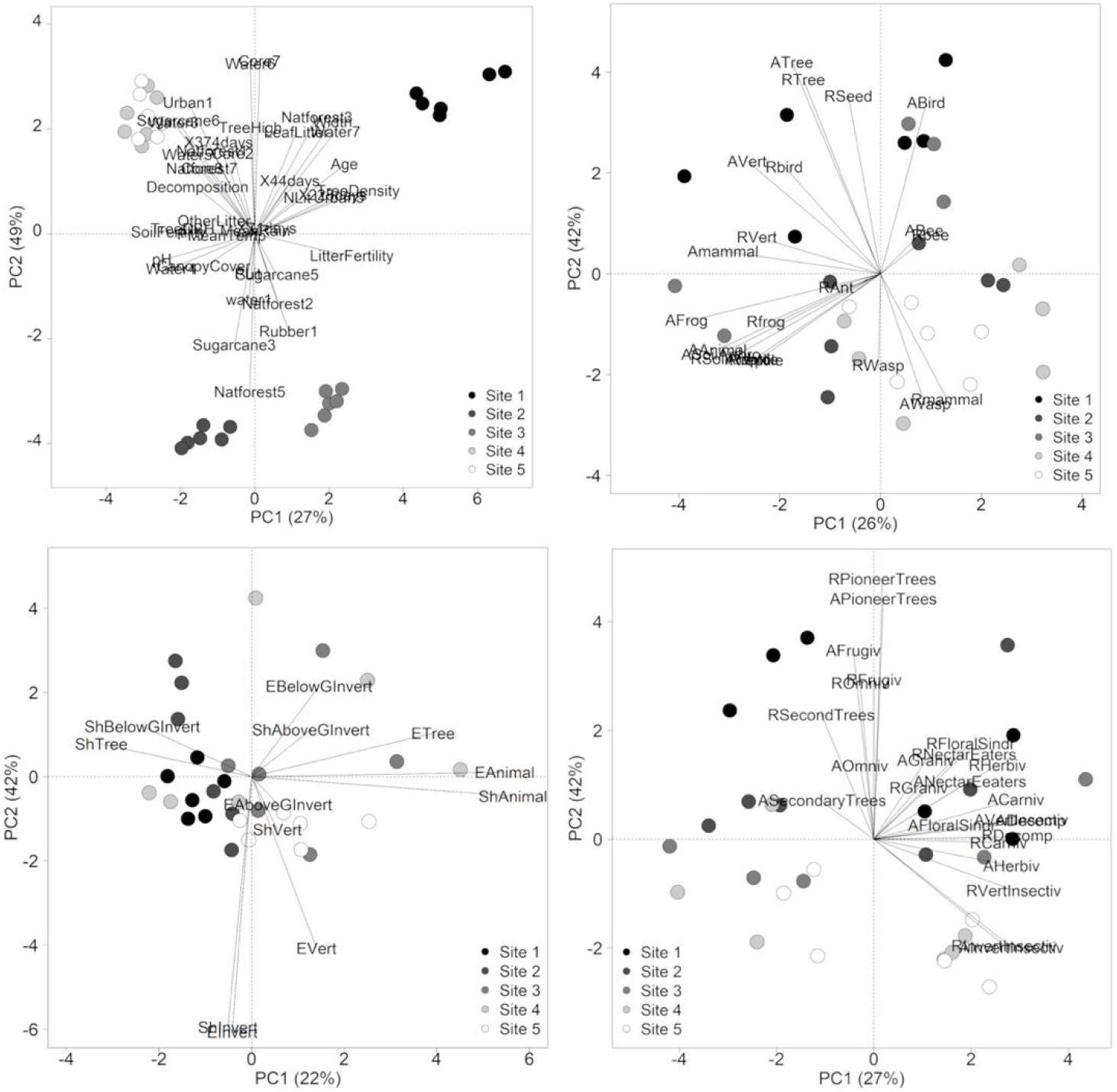
Representation of the first two principal components showing sites dissimilarities according to (a) ecosystem functions and predictors variables at local and landscape levels (see hierarchies in Figure 1); (b) Richness and abundance of distinct groups (biodiversity level); (c) Shannon and Evenness diversity indexes of distinct groups (biodiversity level) and; (d) Richness and abundance of functional groups (biodiversity level) (A = abundance, Arthro = Arthropods, Carniv = Carnivores, E = Evenness Index, Frugiv = Frugivores, G = Ground, Graniv = Granivores, Herbiv = Herbivores, Invert = Invertebrates, Ominv = Omnivores, R = Richness, Second = Secondary, Sh = Shannon Diversity Index, Sindr = Syndrome, Vert = Vertebrates).

**Supplementary Figure 2:**
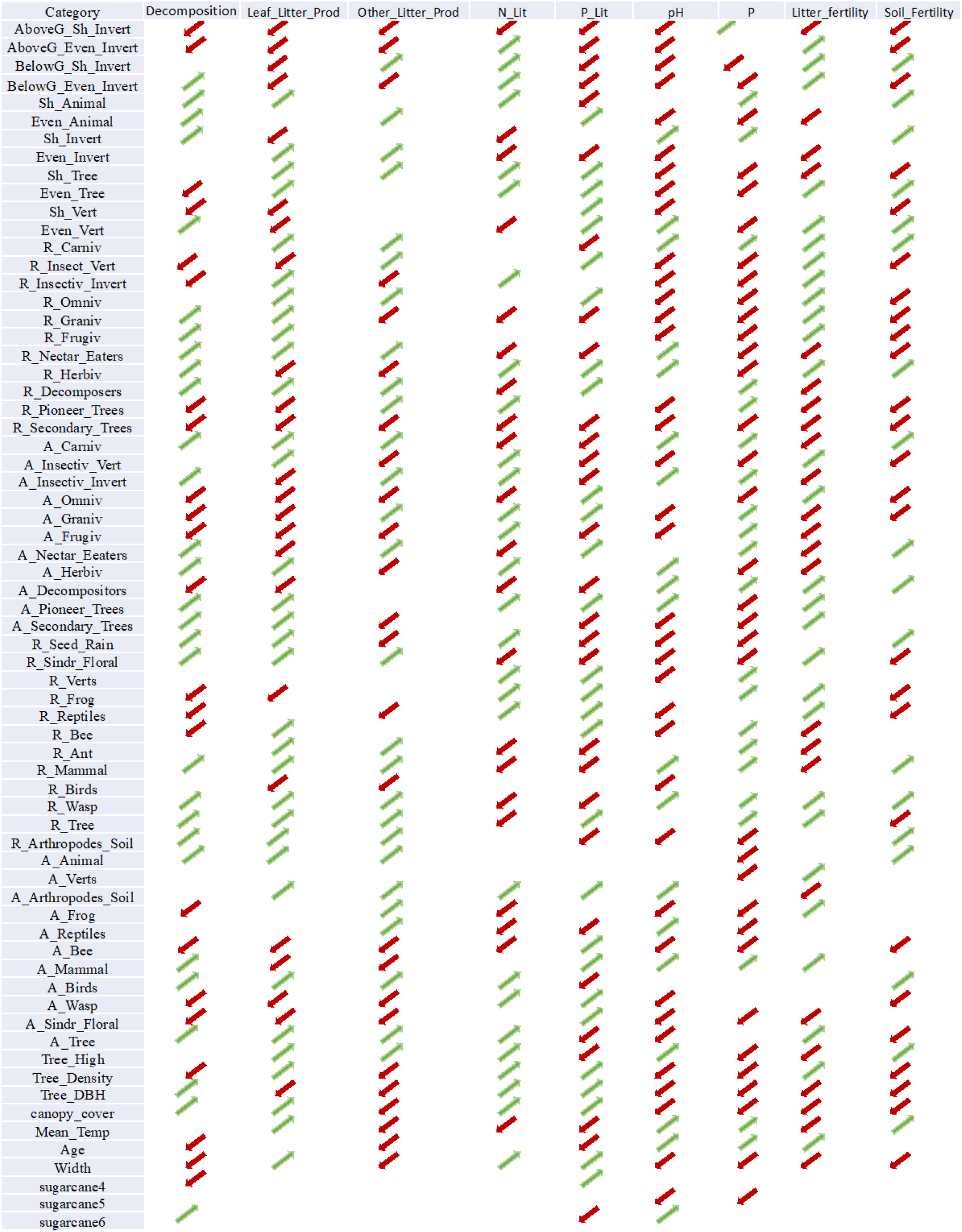
Graphic representation of the positive effects (green arrow pointing northeast) and negative effects (red arrows pointing southwest) of biodiversity, local and landscape variables on the ecosystem functions in a restored area in Southeastern Brazil.

## Suplementary Methods

### Methods

The study was conducted in five patches of riparian forest (hereafter referred to as sampling units) in the region of the reservoir of the Volta Grande hydroelectric power plant (HPP) located on the Rio Grande river that forms the border between Minas Gerais and São Paulo states, Brazil (20°01′54″ S / 48°13′17″ W) (Table S1).

**Supplementary Figure 3:**
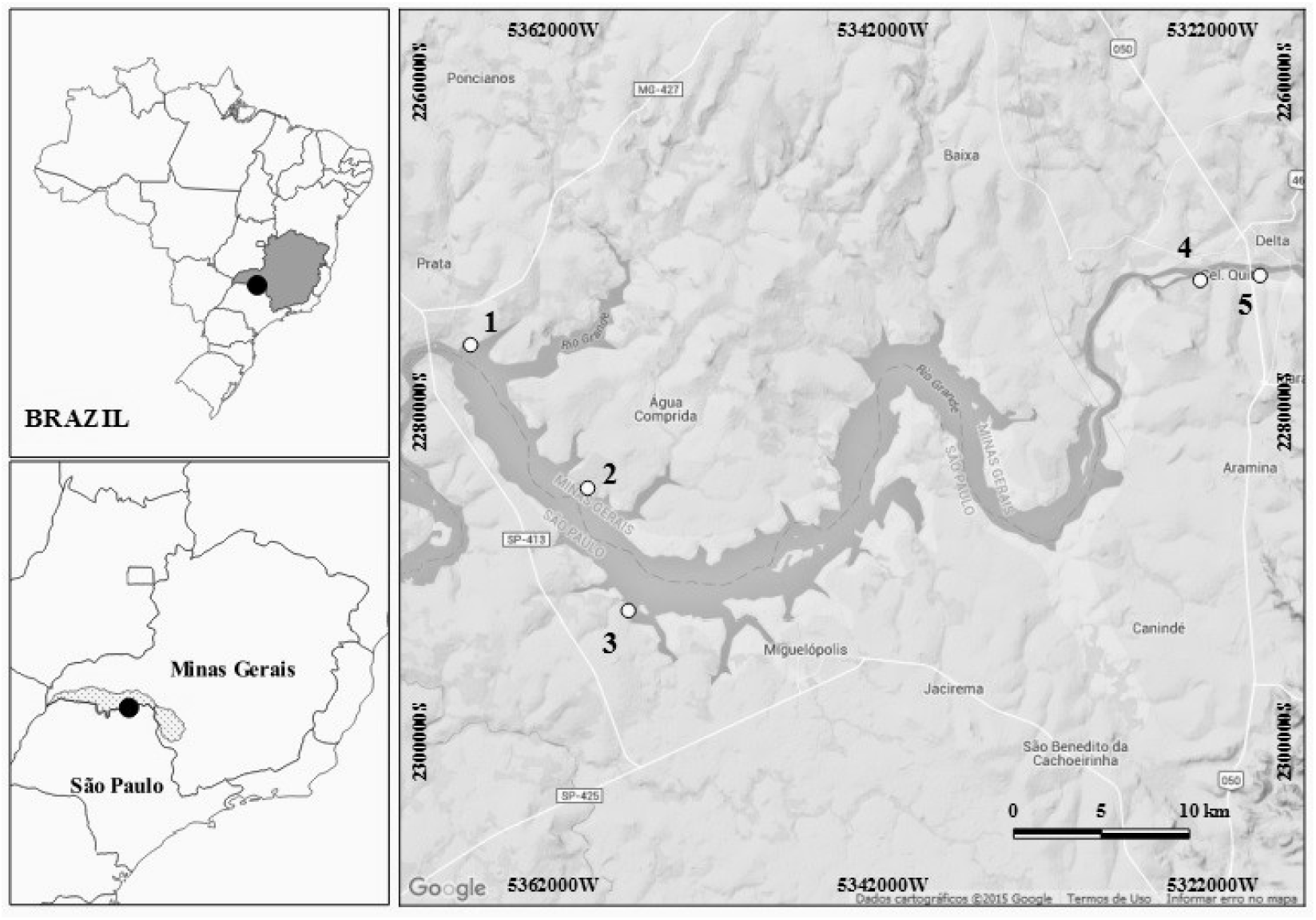
Grande River basin, located in the border of Minas Gerais and São Paulo States, southeastern Brazil. The five riparian forest patches are shown, being areas 1 and 2 located in Minas Gerais State and areas 3, 4 and 5 located in São Paulo State.

**Supplementary Table 1:** Location (UTM) and characteristics of the five riparian forest fragments studied.

| Area | Vegetation | UTM | Location | Municipality | Age (y) | Width (m) |
| --- | --- | --- | --- | --- | --- | --- |
| 1 | Native | 22k | 791531/7783262 | Conceição das Alagoas/MG | 30 | 400 |
| 2 | Reforestation | 22k | 798082/7775015 | Água Comprida/MG | 10 | 30 |
| 3 | Reforestation | 22k | 800294/7768027 | Miguelópolis/SP | 20 | 30 |
| 4 | Reforestation | 23k | 205429/7786874 | Igarapava/SP | 10 | 100 |
| 5 | Reforestation | 23k | 208838/7787209 | Igarapava/SP | 20 | 100 |

**Supplementary table 2.** Units for each variable used in the models

| Variable | Unit |
| --- | --- |
| Richness | S |
| Abundance | N |
| Tree_High | meters |
| Tree_Density | n individuals/m<sup>2</sup> |
| Tree_DBH | cm |
| Decomposition | Remaining mass (X1) |
| Leaf_Litter_Prod | mg ha<sup>-1</sup> |
| Other_Litter_Prod | mg ha<sup>-1</sup> |
| N_Lit | Concentration |
| P_Lit | Concentration |
| pH | absolute value |
| P | Concentration |
| Litterfertility_Index | True value |
| SoilFertility_index | True value |
| Age | Years |
| Width | Meters |

## Plant sampling

In each plot floristic analysis was used to obtain the richness and abundance of trees with diameter at breast height or 1.30 m (DBH) > 10 cm. To describe tree community structure, the traditional quantitative parameters proposed by Mueller-Dombois and Ellenberg (1974) were calculated per plot: absolute density (AD. ind/m2); three high (m) and DBH (cm). Vouchers of the plants studied can be found in the herbarium OUPR, Ouro Preto, Minas Gerais. Brazil.

## Syndromes of pollination

Among the plants with DBH < 10 cm, only flowering plants were recorded for identification of the pollination syndrome. All flowering plants found in the plots were classified by their syndromes, following Faegri & van der Pijl (1979) including lianas, shrubs and herbaceous plants.

## Seed rain and seedling settlement

Seed rain was sampled using mesh with PVC support. This commonly used trap design in tropical rain forests consists of a frame of PVC tubing that supports a net (Harms et al. 2000. Muller-Landau et al. 2002. Wright et al. 1999). We used a trap with a collection area of 1 m2 with a polyester net bag, with holes of less than 1 mm, which was supported by 4 PVC tubes at 0.8 m from the ground. For each sampling site were installed four sampling plots with 3 sampling bags in each plot. Each sampling plot was separated by 100 metros distance. Seed rain was sampled monthly. Seeds collected in traps were counted and identified to the lowest taxonomic level possible to obtain values of richness and abundance.

## Vertebrate sampling

### Small mammal’s, amphibians and reptiles

Pitfall traps were used to collect small mammals (Corn 1994; Cechin and Martins 2000), amphibians (frogs) and reptiles (lizards and snakes). Thirty 60-1 buckets were disposed in three linear transects of 50m; 10 buckets, 5m equidistant, were disposed in each transect. Pitfall traps stood opened for four nights per month in each area. Nine months of field work were carried out (March 2013 to January 2014), being four months during the dry season and five months during the rainy season.

Capture and individual manipulation were carried out according to the guidelines of the Commitee of Animal Care and Use of the American Society of Mammalogists (Sikes et al. 2011). Some specimens were collected, sacrificed and deposited in the vertebrate collection of the Laboratory of Vertebrate Zoology of the Federal University of OuroPreto (LZV-UFOP). The study was approved by the Brazilian System of Authorization and Information on Biodiversity (SISBIO license n° 37067-1) and by the Animal Ethics Committee of the Federal University of OuroPreto (protocol n° 2012/55). Richness and abundance of each group was used for the construction of the models.

### Birds

Birds were quantitatively (point count method; Vielliard et al. 2010) and qualitatively (ad libitum observations) sampled in the five riparian forests studied sites. Each site was sampled once per month. from April 2013 to January 2014.

Three points, separated by 350 m, were set in each riparian forest site (Ralph et al. 1995; Vielliard et al. 2010). The researchers remained in each point for 20 minutes, recording all bird species that were observed or heard. The sampling order was determined using the Latin square method (Bailey 1996) and the surveys occurred only in the mornings (between 06:00 to 09:00 h). Ad libitum observations were conducted in each site between 06:00 to 11:00 and 13:00 to 18:00 hours, resulting in a sampling effort of 10h/day (400 h in total for all five sites). Piacintini et al. (2015) was used for birds’ nomenclature. Birds were identified visually with binoculars, by photography (Canon PowerShot SX50 HS) or audibly. A Sony ICDPX312 recorder was used to document vocalizations.

### Invertebrates sampling

Soil Invertebrate Pitfall traps were also used to sample terrestrial invertebrates (Holway 2005). Ten 400ml plastic cups, filled with a mixture of alcohol 70% and detergent, were disposed laterally to the small mammal pitfall trap transects. As done for small mammals sampling, pitfall traps stood opened for four nights per month in each area. Nine months of field work were carried out (March 2013 to January 2014). Samples were analyzed in laboratory, and the richness (morphospecies) and abundance of each invertebrate order was determined. Invertebrate sampling was approved by the Brazilian System of Authorization and Information on Biodiversity (SISBIO license n° 37067-1).

### Ants

In each plot three trees were arbitrarily selected, and the ants were collected using one baited pitfall trap per tree (radius = 7 cm; height = 9 cm). The pitfall traps were tied to the trees as close as possible to the crown. and baited with sardine (DF) or sardine and honey (PP). The pitfall traps remained on the trees for 48 h. Ants were identified to the higher possible taxonomic level using specific taxonomic keys.

### Trap-nesting bees and wasps

In each of the five sampling units, 12 plots of 100 m2 were installed. In each plot were installed two woodblocks placed 1.5 meters high in the most central tree, forming a plot, totaling 90 blocks of trap-nests, and 5.400 nesting sites total. Trap nests were black cardboard tubes inside inserted in holes drilled into wood blocks with a total of 45 holes arranged linearly (Camilo et al 1995). Trap nests were uniform in length (150 mm) but varied in their inner diameters (6–16 mm). Traps nests were horizontally tied to tree trunks. about 1.8 m above the soil surface. Occupied cardboard, those closed with soil or plant materials, indicating completed nest construction (Krombein 1967), were collected and taken to the laboratory. New empty cardboards were used to replace those collected. In the laboratory, cardboards brought from the field were kept in a glass assay tube plugged with a cotton wad were kept in the laboratory at room conditions (ca. 15–25 °C). Insects were identified and deposited with their nest material in the Entomological Collection of the Laboratório de Biodiversidade, of the Universidade Federal de Ouro Preto.

### Functional Biodiversity

Recorded animals were separated into functional biodiversity. The selected functions are related more to feeding habits, or life forms. Vertebrates were classified as insectivores, carnivores, frugivores, omnivores, nectar eaters, granivores. Invertebrates were classified as herbivores, decomposers, insectivores and nectar eaters. Plants were classified as pioneer or secondary. We also classified plants in accordance with the floral syndrome.

### Ecosystem functions

#### Soil content of Nitrogen, Phosphorus and pH

In each plot three samples of soil (0–10 cm deep) were randomly collected and mixed for chemical and granulometric analyses. Soil content of nitrogen (mg/dm3) and phosphorus (mg/dm3), besides pH in H2O was determined following the procedures described by EMBRAPA (1997).

#### Litter and other litter production

Five 50 × 50 cm (0.25 m2) litter-fall traps were spaced in each sampling plot. Traps were placed 1 m above the ground, taking care to avoid placing traps directly below obvious understory obstructions. Trap contents at each site were monthly collected for 12 months and oven-dried at 50 °C for 3 days before being separated into four constituent parts – leaves and others (fruits, flowers, branches) – and weighed to the nearest 0.1 g. From this, we calculated the leaf litter and other production (g.m-2.year-1) per plot.

#### Litter decomposition

Twelve litter bags (representing four bags for each plot) were installed, coinciding with the location of the litter fall traps. Each litter bag was filled with 10 g of a previous homogenized litter fall sampling. Litterbags (10 cm × 10 cm) were made from 2 mm nylon mesh material. The bags were placed in the forest in the beginning of the dry season and one-quarter of the bags were collected within 44, 71, 228 and 374 days. Collected bags were oven-dried at 50 °C and weighed to the nearest 0.1 g after removing (and noting the presence of) fine root matter, termite runs and mud. The oven-dried litter samples were ground and sieved through a 0.5 mm mesh and analyzed for total N and P concentrations (g.kg-1). The remaining mass for each period (X1) was determined and compared to the initial mass values (X0) using the formula: Xt=X0 e-kt. As proposed by Olson (1963) and (Bockheim et al. 1991), the time required for 50% and 95% mass loss and nutrient release was calculated as t50% = 0.693/k and t95% = 3/k. With this was obtained the Coefficient of decomposition (K).

#### Litter and Soil Fertility index

Following Moran et al. (1998; 2000a,b) we used a soil fertility index (SFI) and litter Fertility Index to explore the relationship between soil and litter fertility with biodiversity. For this we used the equation SFI= pH + OM + P + K + Ca + Mg – Al for Soil Fertility and LFI= OM + P + K + Ca + Mg – Al for litter fertility.

## References

Aczel B, Szaszi B, Holcombe AO (2021) A billion-dollar donation: estimating the cost of researchers’ time spent on peer review. Research Integrity and Peer Review, 6, 14. https://doi.org/10.1186/s41073-021-00118-2

Allan, E., Weisser, W. W., Fischer, M., Schulze, E. D., Weigelt, A., Roscher, C., Schmid, B. (2013). A comparison of the strength of biodiversity effects across multiple functions. Oecologia, 173(1), 223–237. https://doi.org/10.1007/s00442-012-2589-0

Allen, D. C. (2016). Microclimate modification by riparian vegetation affects the structure and resource limitation of arthropod communities. Ecosphere, 7(2), 1–12. https://doi.org/10.1002/ecs2.1200

Alvares, C. A., Stape, J. L., Sentelhas, P. C., De Moraes Gonçalves, J. L., & Sparovek, G. (2013). Köppen’s climate classification map for Brazil. Meteorologische Zeitschrift, 22(6), 711–728. https://doi.org/10.1127/0941-2948/2013/0507

Araújo, G. J., Monteiro, G. F., Messias, M. C. T. B., & Antonini, Y. (2018). Restore it, and they will come: trap-nesting bee and wasp communities (Hymenoptera: Aculeata) are recovered by restoration of riparian forests. Journal of Insect Conservation, 22(2), 245–256. https://doi.org/10.1007/s10841-018-0058-8

Balvanera, P., Siddique, S., Dee, L. Paquette, A., Isbell, F., Gonzalez, A., Griffinet, JN. (2013). Linking biodiversity and ecosystem services: current uncertainties and the necessary next steps. BioScience, 65(1), 49–57. https://doi.org/10.1093/biosci/bit003.

Balvanera, P., Pfisterer, A. B., Buchmann, N., He, J. S., Nakashizuka, T., Raffaelli, D., & Schmid, B. (2006). Quantifying the evidence for biodiversity effects on ecosystem functioning and services. Ecology Letters, 9(10), 1146–1156. https://doi.org/10.1111/j.1461-0248.2006.00963.x

Bardgett, R. D., Mommer, L., & De Vries, F. T. (2014). Going underground: Root traits as drivers of ecosystem processes. Trends in Ecology and Evolution, 29(12), 692–699. https://doi.org/10.1016/j.tree.2014.10.006

Brockerhoff, E. G., Barbaro, L., Castagneyrol, B., Forrester, D. I., Gardiner, B., González-Olabarria, J. R., Jactel, H. (2017). Forest biodiversity, ecosystem functioning and the provision of ecosystem services. Biodiversity and Conservation, 26(13), 3005–3035. https://doi.org/10.1007/s10531-017-1453-2

Bunnell, F. L., & Houde, I. (2010). Down wood and biodiversity - Implications to forest practices. Environmental Reviews, 18(1), 397–421. https://doi.org/10.1139/A10-019

Burdon, F. J., Ramberg, E., Sargac, J., Forio, M. A. E., de Saeyer, N., Mutinova, P. T., McKie, B. G. (2020). Assessing the benefits of forested riparian zones: A qualitative index of riparian integrity is positively associated with ecological status in European streams. Water (Switzerland), 12(4). https://doi.org/10.3390/W12041178

Capellesso, E. S., Scrovonski, K., Zanin, E., Hepp, L., Bayer, C., & Sausen, T. L. (2016). Effects of forest structure on litter production, soil chemical composition and litter–soil interactions. Acta Botanica Brasilica, 30(3), 329–335. https://doi.org/10.1590/0102-33062016abb0048

Cardinale, B. J., Duffy, J. E., Gonzalez, A., Hooper, D. U., Perrings, C., Venail, P., Naeem, S. (2012). Biodiversity loss and its impact on humanity. Nature, 486(7401), 59–67. https://doi.org/10.1038/nature11148

Cardinale, B. J., Matulich, K. L., Hooper, D. U., Byrnes, J. E., Duffy, E., Gamfeldt, L., Gonzalez, A. (2011). The functional role of producer diversity in ecosystems. American Journal of Botany, 98(3), 572–592. https://doi.org/10.3732/ajb.1000364

Cardinale, B. J., Srivastava, D. S., Duffy, J. E., Wright, J. P., Downing, A. L., Sankaran, M., Jouseau, C. (2006). Effects of biodiversity on the functioning of trophic groups and ecosystems. Nature, 443, 989–992. https://doi.org/10.1038/nature05202

Casas, G., Darski, B., Ferreira, P. M. A., Kindel, A., Müller, S. C. (2016). Habitat structure influences the diversity, richness and composition of bird assemblages in successional Atlantic rain forests Grasiela. Tropical Conservation Science, 9(1), 503–524. https://doi.org/10.1177/194008291600900126

Cheng, K., & Heidari, Z. (2019). A new method for quantifying cation exchange capacity: Method verification and application to organic-rich Mudrock formations. Applied Clay Science, 181, 1–15. https://doi.org/10.1016/j.clay.2019.105229

Condit, R., Engelbrecht, B. M. J., Pino, D., Pérez, R., Turnera, B. L. (2013). Species distributions in response to individual soil nutrients and seasonal drought across a community of tropical trees. Proceedings of the National Academy of Sciences of the United States of America, 110(13), 5064–5068. https://doi.org/10.1073/pnas.1218042110

Dambros, J., Vindica, V. F., Delabie, J. H. C., Marques, M. I., Battirola, L. D. (2018). Canopy ant assemblage (Hymenoptera: Formicidae) in two vegetation formations in the Northern Brazilian Pantanal. Sociobiology, 65(3), 358–369. https://doi.org/10.13102/sociobiology.v65i3.1932

Dawud, S. M., Raulund-Rasmussen, K., Ratcliffe, S., Domisch, T., Finér, L., Joly, F. X., Vesterdal, L. (2017). Tree species functional group is a more important driver of soil properties than tree species diversity across major European forest types. Functional Ecology, 31(5), 1153–1162. https://doi.org/10.1111/1365-2435.12821

Díaz, S., Lavorel, S., De Bello, F., Quétier, F., Grigulis, K., Robson, T. M. (2007). Incorporating plant functional diversity effects in ecosystem service assessments. Proceedings of the National Academy of Sciences of the United States of America, 104(52), 20684–20689. https://doi.org/10.1073/pnas.070471610

Fahrig, L. (2003). Effects of Habitat Fragmentation on Biodiversity. Annual Review of Ecology, Evolution, and Systematics, 34, 487–515. https://doi.org/10.1146/annurev.ecolsys.34.011802.132419

Fayle, T. M., Turner, E. C., Basset, Y., Ewers, R. M., Reynolds, G., Novotny, V. (2015). Whole-ecosystem experimental manipulations of tropical forests. Trends in Ecology and Evolution, 30(6), 334–346. https://doi.org/10.1016/j.tree.2015.03.010

Foley, J. A., DeFries, R., Asner, G. P., Barford, C., Bonan, G., Carpenter, S. R., Snyder, P. K. (2005). Global consequences of land use. Science, 309, 570–574. https://doi.org/10.1126/science.1111772

Fonte, S. J., Schowalter, T. D. (2005). The influence of a neotropical herbivore (Lamponius portoricensis) on nutrient cycling and soil processes. Oecologia, 146(3), 423–431. https://doi.org/10.1007/s00442-005-0203-4

Fragoso, J. M. V., Huffman, J. M. (2000). Seed-dispersal and seedling recruitment patterns by the last Neotropical megafaunal element in Amazonia, the tapir. Journal of Tropical Ecology, 16(3), 369–385. https://doi.org/10.1017/S0266467400001462

Friedman, J., Hastie, T., Tibshirani, R. (2010). Regularization Paths for Generalized Linear Models via Coordinate Descent. Journal of Statistical Software, 33(1), 1–22.

Fujii, S., Mori, A. S., Koide, D., Makoto, K., Matsuoka, S., Osono, T., Isbell, F. (2017). Disentangling relationships between plant diversity and decomposition processes under forest restoration. Journal of Applied Ecology, 54(1), 80–90. https://doi.org/10.1111/1365-2664.12733

Gardner, T. A., Barlow, J., Chazdon, R., Ewers, R. M., Harvey, C. A., Peres, C. A., Sodhi, N. S. (2009). Prospects for tropical forest biodiversity in a human-modified world. Ecology Letters, 12(6), 561–582. https://doi.org/10.1111/j.1461-0248.2009.01294.x

Hättenschwiler, S., Coq, S., Barantal, S., Handa, I. T. (2018). Leaf traits and decomposition in tropical rainforests: revisiting some commonly held views and towards a new hypothesis. New Phytologist, 189(4), 950–965. https://doi.org/10.1111/j.1469-8137.2010.03483.x

Hector, A., Bagchi, R. (2007). Biodiversity and ecosystem multifunctionality. Nature, 448, 188–190. https://doi.org/10.1038/nature05947

Hobbie, S. E. (2015). Plant species effects on nutrient cycling: revisiting litter feedbacks. Trends in Ecology and Evolution, 30(6), 357–363. https://doi.org/10.1016/j.tree.2015.03.015

Hooper, D.U., Hapin, F. S., Ewel, J. J., Hector, A., Inchausti, P., Lavorel, S., Wardle, D. A. (2005). Fighting sudden oak death with fire? Science, 1(75), 3–35. https://doi.org/10.1126/science.305.5687.1101

Hooper, David U., Adair, E. C., Cardinale, B. J., Byrnes, J. E. K., Hungate, B. A., Matulich, K. L., Connor, M. I. (2012). A global synthesis reveals biodiversity loss as a major driver of ecosystem change. Nature, 486, 105–108. https://doi.org/10.1038/nature11118

Horn, R., Hartmann, A., Gra, W. (1998). Cation exchange processes in structured soils at various hydraulic properties. Science, 47, 67–72.

Houlton, B. Z., Wang, Y. P., Vitousek, P. M., Field, C. B. (2008). A unifying framework for dinitrogen fixation in the terrestrial biosphere. Nature, 454, 327–330. https://doi.org/10.1038/nature07028

Huang, Y., Ma, Y., Zhao, K., Niklaus, P. A., Schmid, B., & He, J. S. (2017). Positive effects of tree species diversity on litterfall quantity and quality along a secondary successional chronosequence in a subtropical forest. Journal of Plant Ecology, 10(1), 28–35. https://doi.org/10.1093/jpe/rtw115

Kerdraon, D., Drewer, J., Chung, A. Y. C., Majalap, N., Slade, E. M., Bréchet, L., Sayer, E. J. (2020). Litter inputs, but not litter diversity, maintain soil processes in degraded tropical forests—A cross-continental comparison. Frontiers in Forests and Global Change, 2, 1–14. https://doi.org/10.3389/ffgc.2019.00090

Kozovits, A. R., Bustamante, M. M. C., Garofalo, C. R., Bucci, S., Franco, A. C., Goldstein, G., & Meinzer, F. C. (2007). Nutrient resorption and patterns of litter production and decomposition in a Neotropical Savanna. Functional Ecology, 21(6), 1034–1043. https://doi.org/10.1111/j.1365-2435.2007.01325.x

Little, C., Cuevas, J. G., Lara, A., Pino, M., & Schoenholtz, S. (2015). Buffer effects of streamside native forests on water provision in watersheds dominated by exotic forest plantations. Ecohydrology, 8(7), 1205–1217. https://doi.org/10.1002/eco.1575

Londe, V., Messias, M. C. T. B., de Sousa, H. C. (2020). Vegetation restoration is associated with increasing forest width. New Forests, 52(1), 129–144. https://doi.org/10.1007/s11056-020-09786-2

Loreau, M., Hector, A. (2001). Partitioning selection and complementarity in biodiversity experiments. Nature, 412(6842), 72–76. https://doi.org/10.1038/35083573

Loreau, MA, Downing, M., Emmerson, A., Gonzalez, J., Hugues, P., Inchausti, J., Sala, O. (2002). A new look at the relationship between diversity and stability. Oxford University Press. 79–91 p.

Maes, J., Paracchini, M. L., Zulian, G., Dunbar, M. B., & Alkemade, R. (2012). Synergies and trade-offs between ecosystem service supply, biodiversity, and habitat conservation status in Europe. Biological Conservation, 155(2012), 1–12. https://doi.org/10.1016/j.biocon.2012.06.016

Mafia, P. de O., de Azevedo, C. S. (2020). Avifauna of the region of the volta grande hydroelectric power plant in Southeast Brazil. Papeis Avulsos de Zoologia, 60, 1–15. https://doi.org/10.11606/1807-0205/2020.60.16

Manning, P., Van Der Plas, F., Soliveres, S., Allan, E., Maestre, F. T., Mace, G., Fischer, M. (2018). Redefining ecosystem multifunctionality. Nature Ecology and Evolution, 2(3), 427–436. https://doi.org/10.1038/s41559-017-0461-7

Margules, C., Pressey, R. (2000). Systematic conservation planning. Nature 405, 243–253 https://doi.org/10.1038/35012251

Martins, A. C. M., Willig, M. R., Presley, S. J., Marinho-Filho, J. (2017). Effects of forest height and vertical complexity on abundance and biodiversity of bats in Amazonia. Forest Ecology and Management, 391, 427–435. https://doi.org/10.1016/j.foreco.2017.02.039

Naeem, Shahid, Duffy, J. E., Zavaleta, E. (2012). The functions of biological diversity in an age of extinction. Science, 336, 1401–1406. https://doi.org/10.1126/science.1215855

Neves, N. S., Feer, F., Salmon, S., CHateil, C., Ponge, J. F. (2010). The impact of red howler monkey latrines on the distribution of main nutrients and on topsoil profiles in a tropical rain forest. Austral Ecology, 35(5), 549–559. https://doi.org/10.1111/j.1442-9993.2009.02066.x

Noss, R. F., Dobson, A. P., Baldwin, R., Beier, P., Davis, C. R., Dellasala, D. A., Tabor, G. (2012). Bolder Thinking for Conservation. Conservation Biology, 26(1), 1–4. https://doi.org/10.1111/j.1523-1739.2011.01738.x

Oelmann, Y., Kreutziger, Y., Temperton, V. M., Buchmann, N., Roscher, C., Schumacher, J., Wilcke, W. (2007). Nitrogen and Phosphorus Budgets in Experimental Grasslands of Variable Diversity. Journal of Environmental Quality, 36(2), 396–407. https://doi.org/10.2134/jeq2006.0217

Oliver, T. H., Isaac, N. J. B., August, T. A., Woodcock, B. A., Roy, D. B., Bullock, J. M. (2015). Declining resilience of ecosystem functions under biodiversity loss. Nature Communications, 6(1), 1–8. https://doi.org/10.1038/ncomms10122

Palmer, M. A., & Filoso, S. (2009). Restoration of ecosystem services for environmental markets. Science, 325, 575–576. https://doi.org/10.1126/science.1172976

Pasut, C., Tang, F. H. M., Maggi, F. (2020). A mechanistic analysis of wetland biogeochemistry in response to temperature, vegetation, and nutrient input changes. Journal of Geophysical Research: Biogeosciences, 125(4), 1–20. https://doi.org/10.1029/2019JG005437

Pedrini, S. Dixon, K.W, Cross, A.T. (2020). International Standards for Native Seeds in Ecological Restoration. 28:3 S213–S303

Pollock, M. M., & Beechie, T. J. (2014). Does riparian forest restoration thinning enhance biodiversity? The ecological importance of large wood. Journal of the American Water Resources Association, 50(3), 543–559. https://doi.org/10.1111/jawr.12206

R Development Core Team (2016) R: A Language and Environment for Statistical Computing. R Foundation for Statistical Computing, Vienna.

Redondo-Brenes, A., Montagnini, F. (2006). Growth, productivity, aboveground biomass, and carbon sequestration of pure and mixed native tree plantations in the Caribbean lowlands of Costa Rica. Forest Ecology and Management, 232(1), 168–178. https://doi.org/10.1016/j.foreco.2006.05.067

Rhoades, P. R., Davis, T. S., Tinkham, W. T., Hoffman, C. M. (2018). Effects of seasonality, forest structure, and understory plant richness on bee community assemblage in a southern rocky mountain mixed conifer forest. Annals of the Entomological Society of America, 111(5), 278–284. https://doi.org/10.1093/aesa/say021

Roscher, C., Schumacher, J., Gubsch, M., Lipowsky, A., Weigelt, A., Buchmann, N., … Schulze, E. D. (2012). Using plant functional traits to explain diversity-productivity relationships. PLoS ONE, 7(5). https://doi.org/10.1371/journal.pone.0036760

Schaik, C. P. Van, Mirmanto, E. (2013). Spatial Variation in the Structure and Litterfall of a Sumatran Rain Forest. Biotropica, 17(3), 196–205.

Scheibe, A., Steffens, C., Seven, J., Jacob, A., Hertel, D., Leuschner, C., & Gleixner, G. (2015). Effects of tree identity dominate over tree diversity on the soil microbial community structure. Soil Biology and Biochemistry, 81, 219–227. https://doi.org/10.1016/j.soilbio.2014.11.020

Schoolmaster, D. R., Zirbel, C. R., Cronin, J. P. (2020). A graphical causal model for resolving species identity effects and biodiversity–ecosystem function correlations. Ecology, 101(8). https://doi.org/10.1002/ecy.3070

Schowalter TD, Fonte SJ, Geaghan J, Wang J (2011) Effects of manipulated herbivore inputs on nutrient flux and decomposition in a tropical rainforest in Puerto Rico. Oecologia 167:1141–1149. https://doi.org/10.1007/s00442-011-2056-3

Souza-Campana, D. R., Silva, R. R., Fernandes, T. T., Silva, O. G. de M., Saad, L. P., & Morini, M. S. de C. (2017). Twigs in the leaf litter as ant habitats in different vegetation habitats in southeastern Brazil. Tropical Conservation Science, 10, 1–12. https://doi.org/10.1177/1940082917710617

Souza, S. R., Veloso, M. D. M., Espírito-Santo, M. M., Silva, J. O., Sánchez-Azofeifa, A., Souza e Brito, B. G., & Fernandes, G. W. (2019). Litterfall dynamics along a successional gradient in a Brazilian tropical dry forest. Forest Ecosystems, 6(1). https://doi.org/10.1186/s40663-019-0194-y

Spain, A. V., Tibbett, M., Ridd, M., & McLaren, T. I. (2018). Phosphorus dynamics in a tropical forest soil restored after strip mining. Plant and Soil, 427, 105–123. https://doi.org/10.1007/s11104-018-3668-8

Stevenson, P. R., Guzmán-Caro, D. C. (2010). Nutrient transport within and between habitats through seed dispersal processes by woolly monkeys in North-Western Amazonia. American Journal of Primatology, 72(11), 992–1003. https://doi.org/10.1002/ajp.20852

Stratford, J. A., Şekercioğlu, Ç. H. A. (2015). Birds in forest ecosystems. In K. S. H. Peh, R. T. Corlett, & Y. Bergeron (Eds.), Routledge Handbook of Forest Ecology (pp. 281–296). https://doi.org/10.4324/9781315818290

Surasinghe, T. D., & Baldwin, R. F. (2015). Importance of riparian forest buffers in conservation of stream biodiversity: Responses to land uses by stream-associated salamanders across two southeastern temperate ecoregions. Journal of Herpetology, 49(1), 83–94. https://doi.org/10.1670/14-003

Sweeney, B. W., Czapka, S. J., & Yerkes, T. (2002). Riparian forest restoration: Increasing success by reducing plant competition and herbivory. Restoration Ecology, 10(2), 392–400. https://doi.org/10.1046/j.1526-100X.2002.02036.x

Szefer, P., Carmona, C. P., Chmel, K., Konečná, M., Libra, M., Molem, K., Lepš, J. (2017). Determinants of litter decomposition rates in a tropical forest: functional traits, phylogeny and ecological succession. Oikos, 126(8), 1101–1111. https://doi.org/10.1111/oik.03670

Temperton, V. M., Mwangi, P. N., Scherer-Lorenzen, M., Schmid, B., Buchmann, N. (2007). Positive interactions between nitrogen-fixing legumes and four different neighbouring species in a biodiversity experiment. Oecologia, 151(2), 190–205. https://doi.org/10.1007/s00442-006-0576-z

Tibshirani, R. (1996). Regression Shrinkage and Selection Via the Lasso. Journal of the Royal Statistical Society: Series B (Methodological), 58(1), 267–288. https://doi.org/10.1111/j.2517-6161.1996.tb02080.x

Tiessen, H. (2015). Phosphorus dynamics in tropical soils. Phosphorus: Agriculture and the Environment, (46), 253–262. https://doi.org/10.2134/agronmonogr46.c8

Tilman, D., Reich, P. B., Knops, J. M. H. (2006). Biodiversity and ecosystem stability in a decade-long grassland experiment. Nature, 441, 629–632. https://doi.org/10.1038/nature04742

Van Ruijven, J., Berendse, F. (2010). Diversity enhances community recovery, but not resistance, after drought. Journal of Ecology, 98(1), 81–86. https://doi.org/10.1111/j.1365-2745.2009.01603.x

Vitousek, P. M., Howarth, R. W. (1991). Nitrogen limitation on land and in the sea: How can it occur? Biogeochemistry, 13(2), 87–115. https://doi.org/10.1007/BF00002772

Vleut, I., Levy-Tacher, S. I., De Boer, W. F., Galindo-González, J., Ramírez-Marcial, N. (2013). Can a fast-growing early-successional tree (Ochroma pyramidale, Malvaceae) accelerate forest succession? Journal of Tropical Ecology, 29(2), 173–180. https://doi.org/10.1017/S0266467413000126

Wandeler, H., Sousa-Silva, R., Ampoorter, E., Bruelheide, H., Carnol, M., Dawud, S. M., Muys, B. (2016). Drivers of earthworm incidence and abundance across European forests. Soil Biology and Biochemistry, 99, 167–178. https://doi.org/10.1016/j.soilbio.2016.05.003

Wu, H., Xiang, W., Ouyang, S., Forrester, D. I., Zhou, B., Chen, L., Peng, C. (2019). Linkage between tree species richness and soil microbial diversity improves phosphorus bioavailability. Functional Ecology, 33(8), 1549–1560. https://doi.org/10.1111/1365-2435.13355

Zeugin, F., Potvin, C., Jansa, J., Scherer-Lorenzen, M. (2010). Is tree diversity an important driver for phosphorus and nitrogen acquisition of a young tropical plantation? Forest Ecology and Management, 260(9), 1424–1433. https://doi.org/10.1016/j.foreco.2010.07.020

## References

Moran EF, Brondízio ES, Mausel PW, Wu Y. 1994. Integrating Amazonian vegetation, land use, and satellite data. Bioscience 44: 329–338.

Moran EF, Brondízio ES, Tucker JM, Da Silva-Forsberg MC, Falesi I, McCracken SD. 2000a. Strategies for Amazonian forest restoration: evidence for afforestation in five regions of the Brazilian Amazon. Amazonia at the Crossroads: The Challenge of Sustainable Development, Hall A (ed.). Institute for Latin American Studies, University of London: London; 129–149.

Moran EF, Brondízio ES, Tucker JM, Da Silva-Forsberg MC, McCracken SD, Falesi I. 2000b. Effects of soil fertility and land use on forest succession in Amazonia. Forest Ecology and Management 139: 93–108

